# Nanopore sequencing of human associated methicillin-resistant *Staphylococcus aureus* RWP1 genome for inferring antibiotic resistant gene, plasmid and prophage information

**DOI:** 10.1101/2022.10.05.510945

**Authors:** Zarrin Basharat

## Abstract

Methicillin-resistant *Staphylococcus aureus* (MRSA) is a gram-positive bacterium and responsible for several infections in the human. Here, Oxford nanopore mediated whole genome sequencing is reported for human host associated strain. The genome size obtained was of ~3 MB, while GC content was 32.9%. Approximately 3600 CDSs, 62 tRNAs and 16 rRNAs were identified. Phage segments were present, with two complete intact phages. Type I and type IV restriction elements were also detected, along with five plasmids. Several antibiotic resistance genes were also mined, with a key mutation in the fluoroquinolone resistant *gyrA*, phosphonic acid resistant *murA* transferase and fusidic acid resistant *fusA* genes. Sequence type was determined as 4803. Comparative analysis with MRSA isolates from Pakistan (n=20) revealed an open pan-genome, with 1,934 CDSs forming core genome.

---

Nanopore is increasingly being used for onsite studies and sequencing in outbreaks due to its portability and ease of use. This technology was therefore, utilized for the sequence determination of MRSA isolate RWP1. MRSA is easily transmissible and becoming a public health threat due to emergence of antibiotic resistance (Verma et al., 2021). It is also difficult to resolve as it may persist due to biofilm formation (Carrera-Salinas et al., 2022). A MRSA strain RWP1 was obtained from a hospital laboratory in Rawalpindi, Pakistan and cultured for 24 hours. DNA was extracted using QIAamp DNA Mini Kit (Qiagen®) from the 24 hr culture. The amount of DNA was measured with the Qubit® 2.0 Fluorometer (Invitrogen), according to the supplier’s instructions. Library was prepared with 1 μg DNA according to the prescribed instructions (using kit# SQK-LSK109) https://community.nanoporetech.com/protocols/gDNA-sqk-lsk109). Library was loaded onto nanopore device and obtained bases were called using guppy. After initial data generation, base calling was done. Assembly was done using Flye (Kolmogorov et al., 2020, Kolmogorov et al., 2019, Lin et al., 2016) and annotation with Prokka (Seemann, 2014). Plasmid sequence was inferred using PlasmidFinder-2.0 (https://cge.cbs.dtu.dk/services/PlasmidFinder/). MLST, Restriction element finder, plasmid finder, mobile genetic elements and virulence was done using the Center for Genomic Epidemiology website (https://cge.cbs.dtu.dk/). The conjugative regions of the self-transmissible MGE, including origin of transfer site (oriT), relaxase gene, gene encoding type IV coupling protein (T4CP) and gene cluster for bacterial type IV secretion system (T4SS) were found using OriTFinder (https://tool-mml.sjtu.edu.cn/oriTfinder/oriTfinder.html). Active phage regions were mapped using PHASTER (Arndt et al., 2016). Antimicrobial resistance genes were predicted using the Comprehensive Antibiotic Resistance Database (CARD) (https://card.mcmaster.ca/home).

For comparative analysis with other MRSA isolates of Pakistani origin, pan-genome analysis was done using the BPGA software (Chaudhari et al., 2016), according to previously described parameters (Basharat et al., 2018).

A 2.99 or ~3 MB genome was assembled (Fig. 1A). Type I and type IV restriction enzymes were found. Three out of the five enzymes were found to be methyl directed (Supplementary Table 1). Phaster found two intact phages (Supplementary Table 2). One was 92.2Kb and matched most with a Staphylococcal phage phi2958PVL, having 115 CDSs. The other one was of 94.9 Kb in size and coded for 127 proteins. Origin of transfer mapping was done for looking at composition of the bacterial mobile genetic elements and apart from type IV secretion system genes, type IV coupling protein, relaxase, cell surface elastin binding protein, catalase and tetracycline resistant genes were found to be acquired (Fig. 1B). In total, 75 mobile genetic elements were found on plasmids. These included resistance genes, virulence genes, insertion sequences and transposons. Resistance genes having mobility tendency were *dfrG, blaZ, tet(M)* and *mecA*. While *mecA* is used for identifying the MRSA and usually limited to *S. aureus*, *dfrG* has been previously associated with trimethoprim resistance in African strains of *S. aureus* (Nurjadi et al., 2014). It also appeared in strain causing subclinical bovine mastitis in Brazil (Haubert et al., 2017). In a survey comprising data from 13 years from Holland, this gene, along with blaZ plasmid was present in majority of the SCC*mec* subtype IVc(2B) (Witteveen et al., 2022). It has also been reported from *Listeria innocua* (Li et al., 2021a). *tet(M)* with variety of integrative conjugative elements has been reported in several species other than *S. aureus* (Chalker et al., 2021, Marathe et al., 2021, Li et al., 2021b). Virulence genes showing mobility potential included genes coding for staphylokinase, gamma-hemolysin components, leucocidin D/E, serine protease splA, sp1B, sp1E and aurolysin. Among several type of transposons found, transposon Tn6009 is linked with *mer* operon, involved in mercury resistance (Soge et al., 2008). It is interesting to note that several metal resistance genes are found alongside the antibiotic resistance genes, possibly due to shared mechanism of purging elements harmful to bacteria, out of the organism. All the rest of insertion sequences belonged to either IS256 or IS6 family. Several copies of insertion element IS256 (part of Tn4001 transposon family) were present and known to be linked with gentamicin resistance and biofilm formation (Kozitskaya et al., 2004). Among the twenty MRSA isolates reported from Pakistan and associated with human host, 1,934 genes were shared with the strain RWP1 (Supplementary Table. 3).

**Fig. 1.**
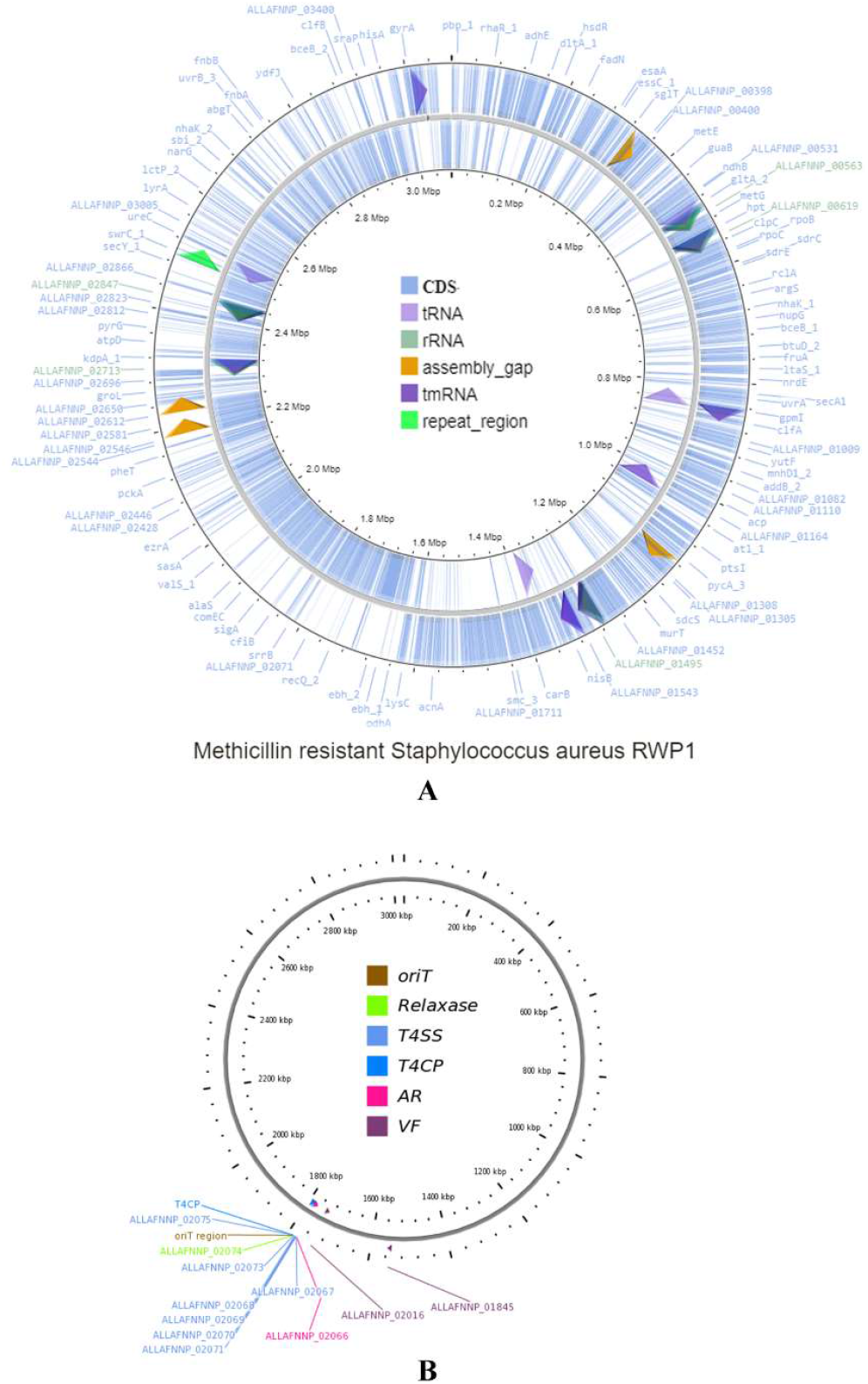
(A) Circular view of the assembled genome. (B) Origin of transfer region showing nearby elements, featuring a *tetR* in pink.

**Table 1.**
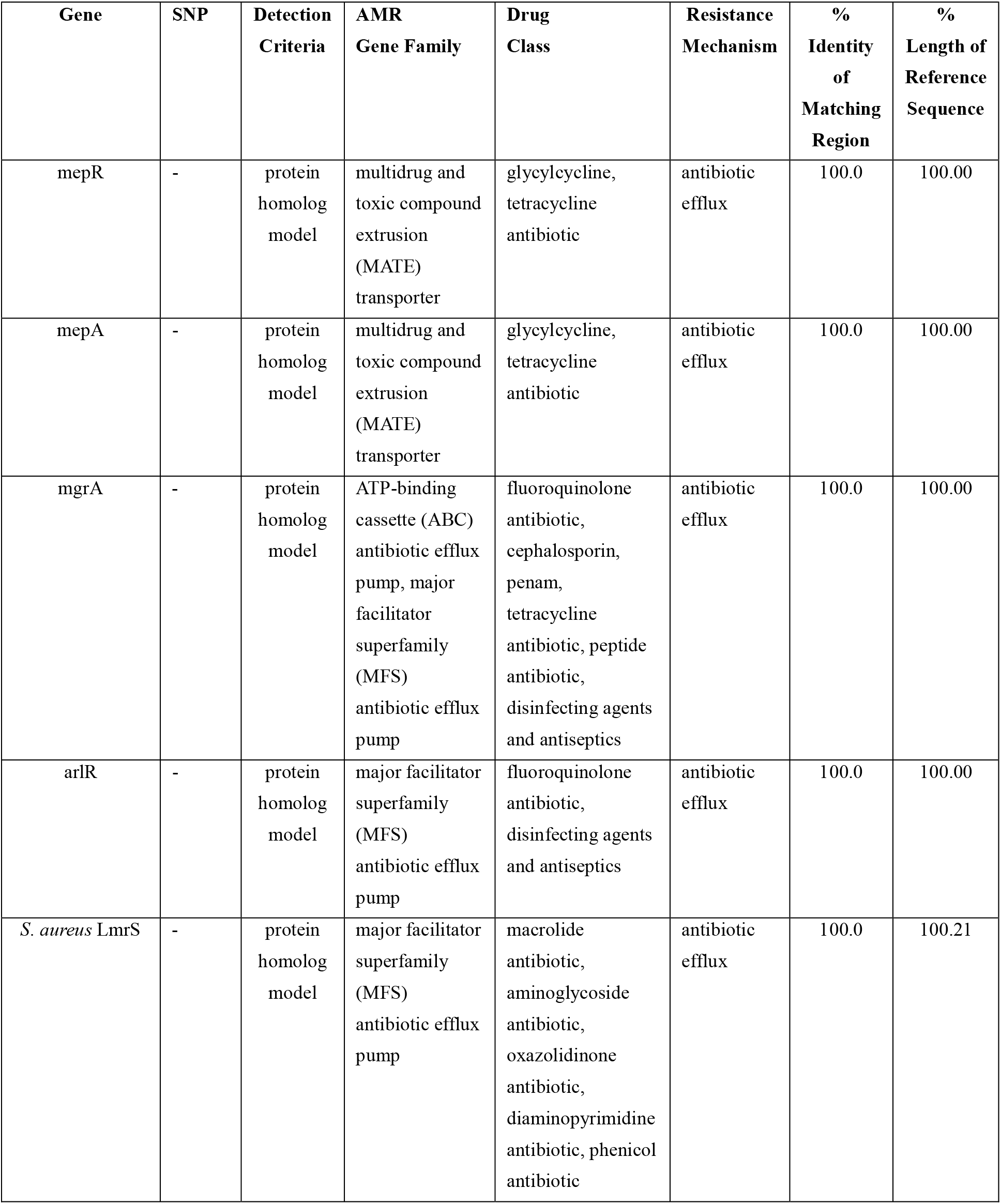
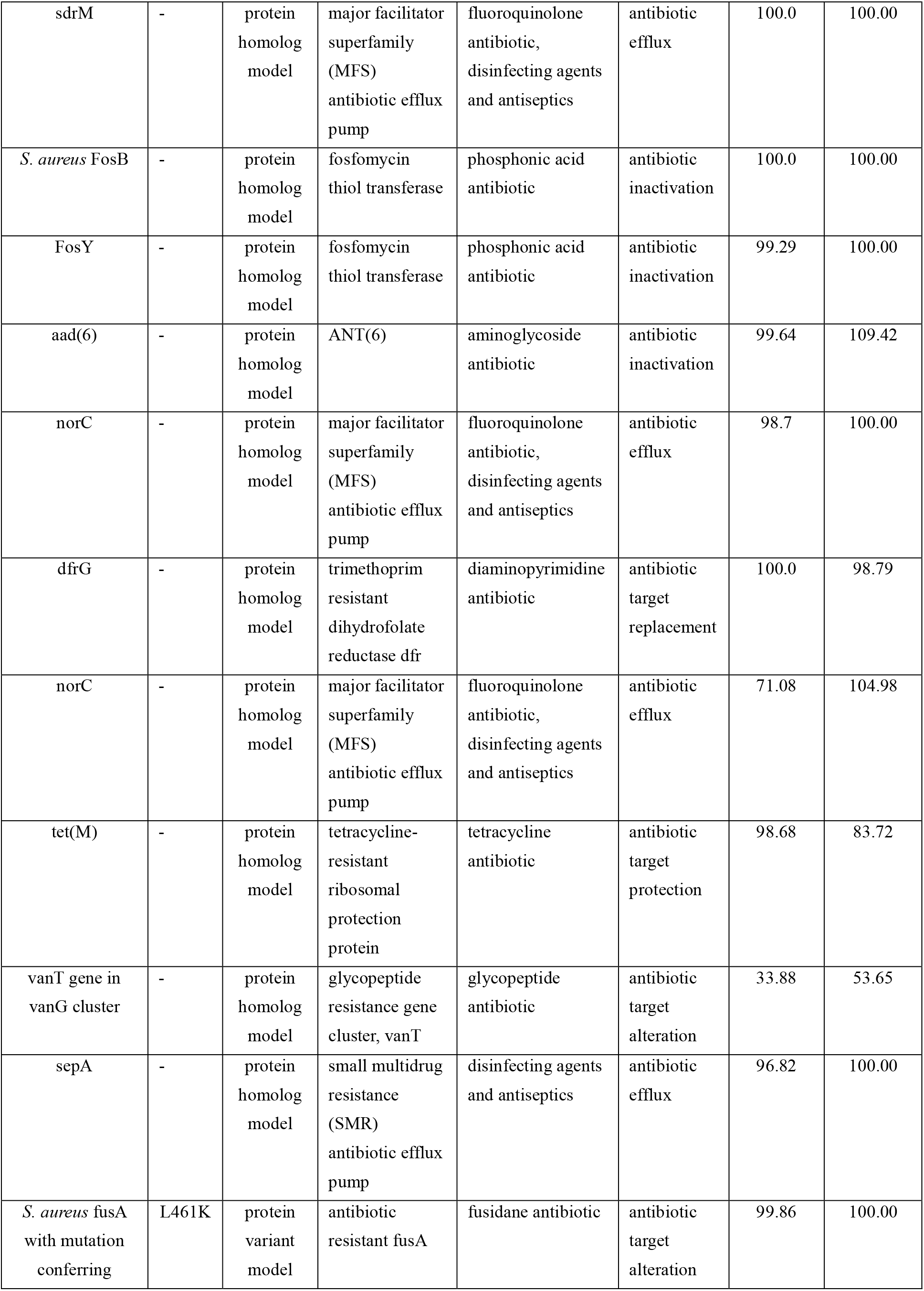
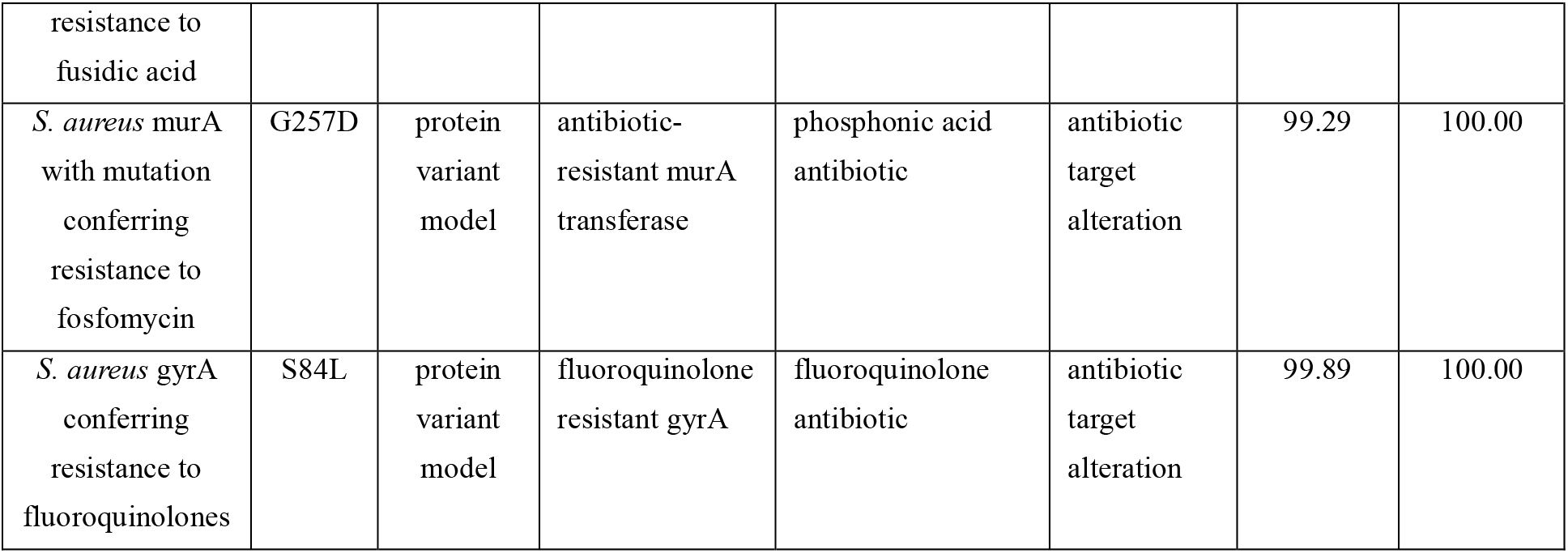
Antibiotic resistance genes in the sequenced MRSA isolate.

## Data availability

The genome has been submitted to NCBI, BioProject ID: PRJNA885976, Biosample ID:SAMN31113469.

