## Supplementary data for "Nanopore sequencing of human associated methicillin-resistant *Staphylococcus aureus* RWP1 genome for inferring antibiotic resistant gene, plasmid and prophage information"

**Supplementary Table 1. Restriction enzymes in MRSA RWP-1.**

| **Type I Restriction enzymes** | | | | | | | | |
| --- | --- | --- | --- | --- | --- | --- | --- | --- |
| **Gene** | **%Identity** | **HSP/Query length** | **Contig** | **Position in contig** | **Type** | **Function** | **Recognition Seq** | **Accession number** |
| *M.SauTCHI* | 99.94 | 1557 / 1557 | Scaffold_1 | 1346769..1348325 | Type I | methyltransferase | CCAYNNNNNNTGT | CP000730 |
| *S.Sau300II* | 99.92 | 1200 / 1200 | Scaffold_1 | 1348318..1349516 | Type I | specificity subunit | CCAYNNNNNNTGT | CP007176 |
| *M.Sau20231I* | 100.00 | 1557 / 1557 | Scaffold_1 | 449948..451504 | Type I | methyltransferase | AGGNNNNNGAT | CP011526 |
| *S.Sau8532II* | 100.00 | 1212 / 1212 | Scaffold_1 | 451497..452708 | Type I | specificity subunit | AGGNNNNNGAT | LN831049 |
| **Type IV Restriction enzymes** | | | | | | | | |
| **Gene** | **%Identity** | **HSP/Query length** | **Contig** | **Position in contig** | **Type** | **Function** | **Recognition Seq** | **Accession number** |
| *SauUSI* | 100.00 | 2862 / 2862 | Scaffold_1 | 2732450..2735311 | Type IV | methyl-directed restriction enzyme | SCNGS | CP000255 |

**Supplementary Table 2. Prophages of MRSA RWP-1.**

Total nine prophage regions were identified, of which two were intact, three were incomplete, four regions were questionable.

| **Region** | **Region_length** | **Completeness** | **Score** | **#cds** | **Region_position** | **Possible phage** | **Gc_percentage** |
| --- | --- | --- | --- | --- | --- | --- | --- |
| [1](http://phast.wishartlab.com/cgi-bin/change_detail_html.cgi?num=1664617491_1#1) | 34.3Kb | questionable | 70 | 35 | [32591-66898](http://phast.wishartlab.com/cgi-bin/get_region_DNA.cgi?num=1664617491_1&number=1) | PHAGE_Staphy_SPbeta_like_NC_029119 | 30.91% |
| [2](http://phast.wishartlab.com/cgi-bin/change_detail_html.cgi?num=1664617491_1#2) | 37.2Kb | questionable | 70 | 51 | [328009-365300](http://phast.wishartlab.com/cgi-bin/get_region_DNA.cgi?num=1664617491_1&number=2) | PHAGE_Staphy_77_NC_005356, ...... | 33.66% |
| [3](http://phast.wishartlab.com/cgi-bin/change_detail_html.cgi?num=1664617491_1#3) | 10.4Kb | incomplete | 60 | 15 | [659413-669910](http://phast.wishartlab.com/cgi-bin/get_region_DNA.cgi?num=1664617491_1&number=3) | PHAGE_Staphy_SPbeta_like_NC_029119 | 30.62% |
| [4](http://phast.wishartlab.com/cgi-bin/change_detail_html.cgi?num=1664617491_1#4) | 14.9Kb | questionable | 90 | 22 | [845461-860367](http://phast.wishartlab.com/cgi-bin/get_region_DNA.cgi?num=1664617491_1&number=4) | PHAGE_Staphy_SPbeta_like_NC_029119 | 32.58% |
| [5](http://phast.wishartlab.com/cgi-bin/change_detail_html.cgi?num=1664617491_1#5) | 36.5Kb | incomplete | 60 | 52 | [1117092-1153592](http://phast.wishartlab.com/cgi-bin/get_region_DNA.cgi?num=1664617491_1&number=5) | PHAGE_Staphy_phiN315_NC_004740 | 33.57% |
| [6](http://phast.wishartlab.com/cgi-bin/change_detail_html.cgi?num=1664617491_1#6) | 47.9Kb | questionable | 80 | 78 | [1218685-1266669](http://phast.wishartlab.com/cgi-bin/get_region_DNA.cgi?num=1664617491_1&number=6) | PHAGE_Staphy_53_NC_007049 | 33.83% |
| [7](http://phast.wishartlab.com/cgi-bin/change_detail_html.cgi?num=1664617491_1#7) | 5.9Kb | incomplete | 50 | 10 | [1522967-1528953](http://phast.wishartlab.com/cgi-bin/get_region_DNA.cgi?num=1664617491_1&number=7) | PHAGE_Staphy_SPbeta_like_NC_029119, ...... | 28.13% |
| [8](http://phast.wishartlab.com/cgi-bin/change_detail_html.cgi?num=1664617491_1#8) | 92.2Kb | intact | 150 | 115 | [1736059-1828281](http://phast.wishartlab.com/cgi-bin/get_region_DNA.cgi?num=1664617491_1&number=8) | PHAGE_Staphy_phi2958PVL_NC_011344 | 34.00% |
| [9](http://phast.wishartlab.com/cgi-bin/change_detail_html.cgi?num=1664617491_1#9) | 94.9Kb | intact | 150 | 127 | [2165418-2260414](http://phast.wishartlab.com/cgi-bin/get_region_DNA.cgi?num=1664617491_1&number=9) | PHAGE_Staphy_77_NC_005356 | 33.06% |

**Supplementary Table 3. Pan-genome statistics of human associated MRSA strains from Pakistan.**

| Genome no. | Assembly accession | Organism name | No. of core genes | No. of accessory genes | No. of unique genes | No. of exclusively absent genes |
| --- | --- | --- | --- | --- | --- | --- |
| 1 | GCA_018995405.1 | 16S | 1934 | 861 | 20 | 10 |
| 2 | GCA_018995425.1 | 25S | 1934 | 679 | 8 | 12 |
| 3 | GCA_018995375.1 | 31S | 1934 | 864 | 8 | 3 |
| 4 | GCA_018995365.1 | 33S | 1934 | 946 | 7 | 0 |
| 5 | GCA_018995345.1 | 34S | 1934 | 788 | 2 | 3 |
| 6 | GCA_018995305.1 | 37S | 1934 | 849 | 2 | 0 |
| 7 | GCA_018995225.1 | 39S | 1934 | 810 | 10 | 4 |
| 8 | GCA_018995265.1 | 41S | 1934 | 929 | 1 | 0 |
| 9 | GCA_018995325.1 | 43S | 1934 | 824 | 13 | 2 |
| 10 | GCA_018995165.1 | 48S | 1934 | 666 | 3 | 39 |
| 11 | GCA_018995115.1 | 50S | 1934 | 707 | 107 | 11 |
| 12 | GCA_005706855.2 | Lr2 | 1934 | 693 | 73 | 9 |
| 13 | GCA_005707335.1 | Lr3 | 1934 | 630 | 19 | 10 |
| 14 | GCA_005705815.1 | Lr6 | 1934 | 689 | 16 | 2 |
| 15 | GCA_005707215.1 | Lr12 | 1934 | 674 | 65 | 13 |
| 16 | GCA_021460465.1 | MIN-169 | 1934 | 689 | 42 | 4 |
| 17 | GCA_021460375.1 | MIN-170 | 1934 | 809 | 5 | 28 |
| 18 | GCA_005706035.2 | P10 | 1934 | 875 | 42 | 3 |
| 19 | GCA_005705635.2 | R46 | 1934 | 762 | 7 | 3 |
| 20 | GCA_005707015.1 | R50 | 1934 | 751 | 40 | 0 |
| **21** | **-** | **RWP1** | **1934** | **718** | **100** | **70** |
